# Fine-Tuning GBS Data with Comparison of Reference and Mock Genome Approaches for Advancing Genomic Selection in Less Studied Farmed Species

**DOI:** 10.1101/2023.10.03.560633

**Authors:** Daniel Fischer, Miika Tapio, Oliver Bitz, Terhi Iso-Touru, Antti Kause, Ilma Tapio

**Author notes:** Corresponding author: Daniel Fischer.

## Abstract

**Background:** Diversifying animal cultivation demands efficient genotyping for enabling genomic selection, but non-model species lack efficient genotyping solutions. The aim of this study was to optimize a genotyping-by-sequencing (GBS) double-digest RAD-sequencing (ddRAD) pipeline. Bovine data was used to automate the bioinformatic analysis. The application of the optimization was demonstrated on non-model European whitefish data.

**Results:** DdRAD data generation was designed for a reliable estimation of relatedness and is scalable to up to 384 samples. The GBS sequencing yielded approximately one million reads for each of the around 100 assessed samples. Optimizing various strategies to create a de-novo reference genome for variant calling (mock reference) showed that using three samples outperformed other building strategies with single or very large number of samples. Adjustments to most pipeline tuning parameters had limited impact on high-quality data, except for the identity criterion for merging mock reference genome clusters. For each species, over 15k GBS variants based on the mock reference were obtained and showed comparable results with the ones called using an existing reference genome. Repeatability analysis showed high concordance over replicates, particularly in bovine while in European whitefish data repeatability did not exceed earlier observations.

**Conclusions:** The proposed cost-effective ddRAD strategy, coupled with an efficient bioinformatics workflow, enables broad adoption of ddRAD GBS across diverse farmed species. While beneficial, a reference genome is not obligatory. The integration of Snakemake streamlines the pipeline usage on computer clusters and supports customization. This user-friendly solution facilitates genotyping for both model and non-model species.

## Background

Humans have successfully domesticated over five hundred animal species, and the number of newly cultivated species has been increasing by at least ten species per year [1,2]. Particularly in recently domesticated species, our understanding of their genetic diversity and the genetic basis of traits may be insufficient. Genome wide data and genomic selection have revolutionized animal breeding by improving productivity [3–5], as well as incorporating health and welfare traits [6,7]. In genomic selection, thousands of DNA markers are used to predict the genomic breeding value of an individual [8,9], but genotyping presents a significant challenge for rare or novel production species. A recent review of genome data [10] revealed that nearly half of the aquaculture species, with an annual production exceeding 350 million kg [11], lack reference genome information, which together with genetic polymorphism characterization is a necessary resource for the development of commercial SNP-chip platforms or targeted genotyping-by-sequencing solutions. Therefore, it is crucial to make cost-effective and reliable alternative genotyping methods widely available for non-model organisms to advance genomic selection and stock management in niche production species.

The advantage of genome-assisted breeding value estimation largely stems from reliable estimation of relationships [12] and a common genomic selection approach is directly based on the genomic relationship matrix (GRM), which estimates the proportion of the genome shared identical by descent between pairs of individuals. This method does not require a genomic map or a reference genome and performs well even with low marker densities (10 SNPs per morgan) [13]. However, additional markers are beneficial and, for example, in Atlantic salmon, densities up to 50 to 200 markers per morgan (1 000 to 5 000 markers in total) have been recommended [4,14]. The accuracy and cost-effectiveness of genomic selection depend on the balance between the number of genotyped markers and individuals, with marker numbers of 1 000 to 2 000 SNPs being suggested [15].

Genotyping-by-sequencing (GBS) [16] is a cost-effective approach for simultaneous genome-wide SNP discovery and genotyping without prior knowledge of the genome sequence. Restriction-site associated DNA sequencing (RAD) [17–19] and double-digest RAD-sequencing (ddRAD) [20,21] are reduced-representation genome sequencing methods that target a small portion of the genome using restriction enzymes. These methods can generate sequencing-libraries from hundreds to hundreds of thousands of fragments genome wide. Both wet lab protocols and parameters used in post-sequencing analysis impact the number of recovered reads, mean sequencing target coverage, recovered genetic loci/marker, and genotype completeness and accuracy [20]. While the number of polymorphic markers is the main concrete criterion for evaluating the suitability of a genotyping method for genomic selection, the actual genotyping goal of reliable estimation of relatedness might be influenced by the minor allele frequencies (MAF), codominant or mendelian inheritance and repeatability. GBS variants typically have a lower call rate per sample and repeatability among sample sets compared to SNP arrays. Additionally, genotyping errors, especially allelic dropouts (as false homozygotes), can introduce bias in the relatedness estimates used in genomic selection [22]. However, optimized GBS pipelines can exhibit high consistency with SNP-chip data [23].

Hence, the primary objective of this study was to optimize the GBS method ddRAD and fine-tune the bioinformatic pipeline parameters for processing and controlling of the high-quality SNP data for genomic selection in non-model species. The second objective was to test the repeatability of the data generation. We fine-tuned the bioinformatics pipeline parameters by utilizing dairy cattle GBS and whole-genome resequencing (WGS) data. Following this, we applied the established data processing routines on data generated for European whitefish (*Coregonus lavaretus* L) using the available reference genome of the closest relative *Coregonus supersum* ‘balchen’ [24]. European whitefish is the second most important farmed fish species in Finland [25,26]. It is also a species used in ecological studies and it is known to have undergone widespread phylogeographic structuring and the repeated evolution of distinct ecological ecotypes [27]. The overarching objective was to make the GBS method simpler to use across diverse species, eliminating the need for extensive bioinformatics expertise or specialized units. This advancement holds the potential to enhance genomic selection and refine animal breeding practices, particularly within less studied species.

## Results

### Restriction enzyme selection in silico

The numbers of double digested genome fragments within the range of 150-400 bp and consequently the expected variant numbers were three to four times more strongly influenced by the choice of the enzyme pair than by the species assessed (Figure 1). The predicted fragment numbers fulfilled the preset criteria for all enzyme pairs, the number of fragments being the lowest for the EcoRI;SphI pair, with approximately 25 – 50 thousand fragments (or 20 – 40 thousand estimated variants). The reference genome based fragment numbers for the two main targets, *Bos taurus (*ARS-UCD1.2), and *Coregonus supersum* (AWG_v2), were for the pair EcoRI;SphI 50 000 and 30 000, for the pair EcoRI;MspI 120 000 and 110 000, for the pair MluCI;SphI 270 000 and 230 000, for the pair EcoRI;MseI 380 000 and 180 000, for the pair EcoRI;NlaIII 440 000 and 200 000, respectively. The predicted fragment number for the EcoRI;SphI pair was within the desired range of 10 000 – 100 000 fragments, which was expected to provide a minimum of 5 000 relatedness informative variants. Moreover, this enzyme pair provided the most uniform distribution of fragments across the size range, reducing the size selection lab protocol choice to the decision of window width (Figure 1). The EcoRI;SphI pair was the most optimal for all the currently assessed species.

**Figure 1.**
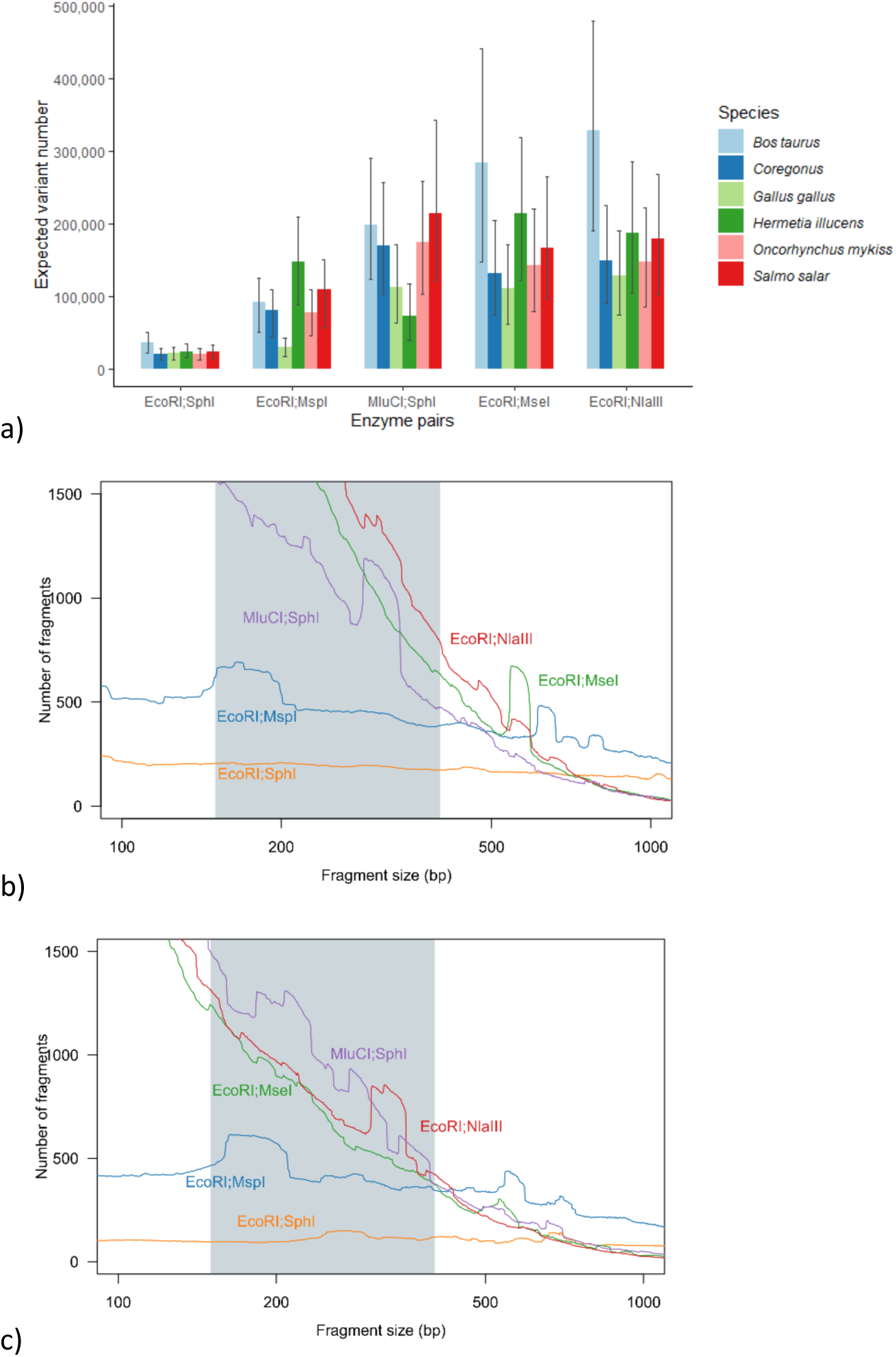
In silico comparison of enzyme pairs. Expected variant numbers across the species and assessed restriction enzyme pairs (a), where whiskers indicate the impact of symmetric widening or narrowing the fragment size range by 100 bp. The predicted frequency distribution of double digested template fragments of different sizes in *Bos taurus* (b) and *Coregonus sp.* (c) averaged over 50 bp window across fragment sizes from 100 to 1 000 bp. The grey area denotes the included size range (150 – 400 bp).

### Raw GBS and WGS sequencing data

GBS sequencing of 36 cow libraries generated in total 43 109 115 PE reads of 2×75 bp in length, with an average of 1 197 475 PE reads per sample. After trimming, 39 730 518 PE reads remained (avg: 1 103 625 reads per sample). Sample details are listed in Table S1. In case of the 66 whitefish libraries sequenced, from the total of 78 577 269 PE reads of 2×75 bp in length (avg: 1 190 565 reads per sample) 71 655 413 reads passed the quality control trimming (avg: 1 086 688 reads per sample). After quality control, the average read length dropped to 66 bp for reads R1 and 60 bp for reads R2.

WGS sequencing of 12 cow samples generated in total 3 918 912 122 PE reads of 2×150 bp in length, with an average of 326 659 344 PE reads per sample. After trimming, 3 865 355 653 PE reads remained with average of 322 112 971 reads per sample.

### GBS fragment recovery

The mapping of the quality-trimmed GBS derived cow data against the non-size selected in-silico (EcoRI;SphI) digested *Bos taurus* (ARS-UCD1.2) reference genome indicated that about 86% of the reads aligned to fragments within the 150 - 400 bp size range (Figure 2). This alignment window was narrower than the expected full insert size range of 150-550 bp. The in-silico digestion simulation generated in total 66 450 genome fragments between 150 and 400 bp in length. Considering that the remaining 14% of the reads were outside this span, our mock reference was expected to have between 66 450 and 79 100 clusters.

**Figure 2:**
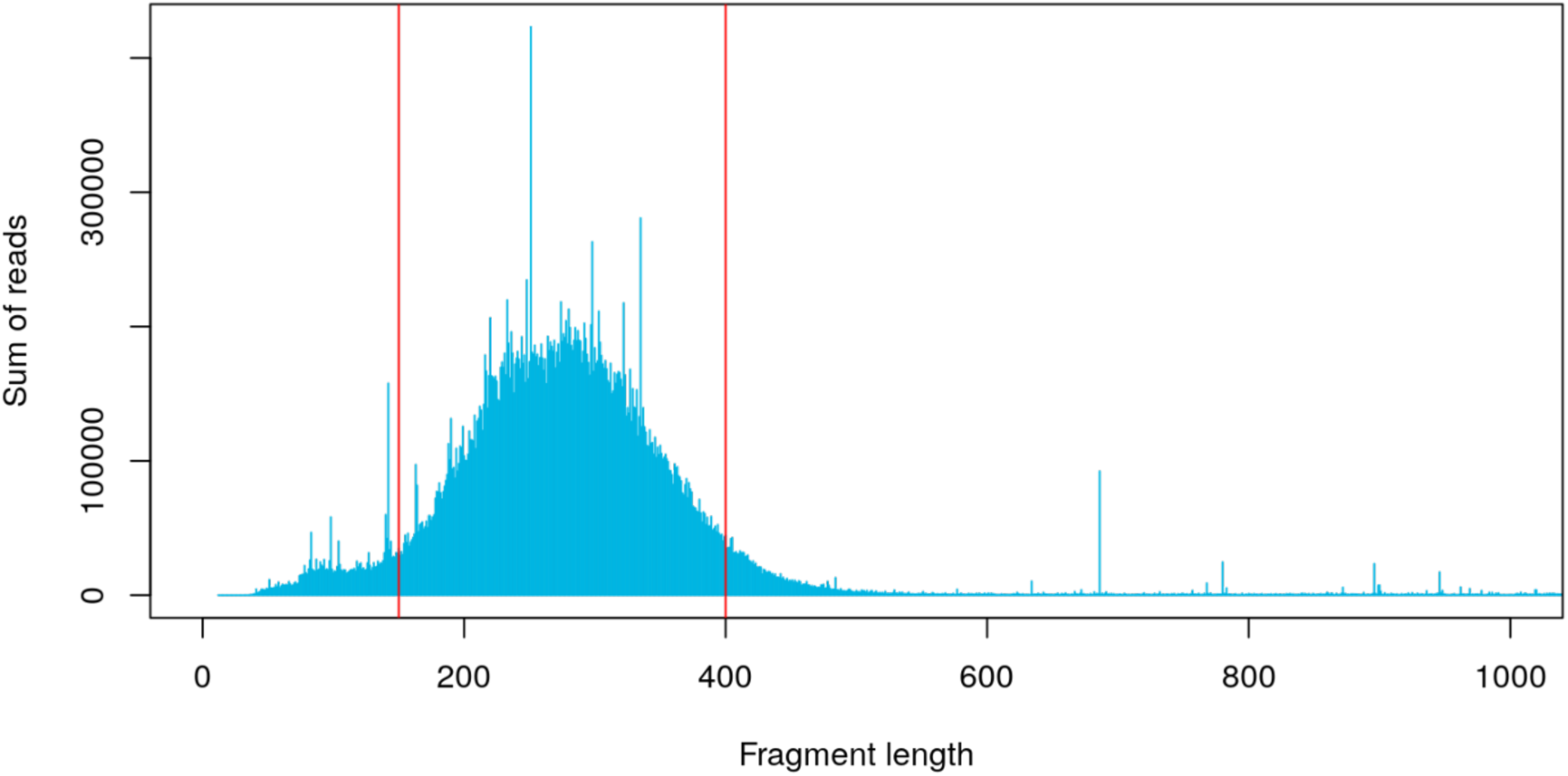
Distribution of quality-trimmed cow GBS reads across in-silico digested *Bos taurus* (ARS-UCD1.2) reference genome fragment lengths. Red vertical lines indicate the boundaries of the estimated effective fragment size.

### Mock reference quality

The construction of a mock reference relies on the defined data and parameter configurations. An evaluation against the size-selected in-silico digested reference, measuring average coverage percentages and secondary alignments (Figure S2), unveiled an over-inflation of the mock reference when utilizing all samples, resulting in the exclusion of mock-strategy 4. While focusing on one sample (mock-strategy 1 and 2) approximated the optimal cluster counts, it introduces the risk of sample-specific biases in the mock reference. As a result, mock-strategy 3 emerged as the preferred choice. However, its advantage over mock-strategy 4 was reduced by the final mock refinement step, which curbed most of the excessive cluster inflation, as indicated by consistent alignment trends nearing the expectation value (Figure S2, gray box).

Adjustments to input data parameters had minimal impact on the mock reference. PE read merging using p-value thresholds (0.001, 0.01, 0.05) yielded consistent mock reference lengths and alignment percentages against the in-silico reference. Around 99.8% of the mock clusters aligned against the reference genome, accompanied by a modest number of unaligned clusters (417-900). Mock cluster counts and secondary alignments remained stable. Parameter pl (min. merged cluster length) showed negligible impact across reasonable values, aligning with expectations. Cluster generation parameters, especially the nucleotide similarity parameter (id), had, however, significant influence. Its extreme values led to drastic changes in the merged cluster numbers, while moderate values (e.g., 0.85) yielded expected alignments. The minimum cluster length (min) and read stitching optimization (rl) parameters had limited impact. Optimal parameters for the mock reference creation were p=0.05, pl=50, id=0.85, min=80 and rl=75 (Figure S3).

For the mock refinement step, strict parameters (e.g., average 10 reads per sample per cluster, ≥10 samples with aligned reads on cluster) appeared optimal for a stable variant set creation. Refined mock references exhibited improved alignment against the *Bos taurus* (ARS-UCD1.2) reference genome (dashed-line), although the average sample-wise alignment of data against the mock reference was slightly decreased for the refined mock compared to pre-refinement mock (Figure S4).

### Variant calling and GBS quality estimation

Applying the GATK best practice variant calling pipeline to the full genome WGS data produced in total 17 376 716 variants for the cow samples, with 42 160 variants intersecting regions on the reference genome that had a minimum coverage of three reads from the GBS data from at least 10 samples.

Aligning GBS data to the reference genome (ARS-UCD1.2) resulted, after similar filtering, in 20 794 variants. Calling variants using the pre-refinement mock reference, based on mock strategy 3, yielded 16 404, and with refinement, 16 416 variants. In the case of GBS, we obtained a MAF of 0.26 (sd: 0.13) using the mock reference and 0.27 (sd: 0.14) while using the reference genome. The average call rate using the GBS approach in combination with the ARS-UCD1.2 reference genome was 94.8%, with average 11.38 (sd: 0.75) samples per variant, respectively 11.37 (sd: 0.76) with using the created mock reference genome. For the WGS, we observed for the 42 160 variants a MAF of 0.21 (sd: 0.14) with a call rate of 99.9% with 11.99 (sd: 0.13) samples having called each variant on average.

The overlap of reference based GBS and WGS variant sets, defined by their chromosomal positions, comprised 18 196 loci, representing approximately 87.5% alignment between the GBS and WGS datasets. These variants exhibited a WGS-based MAF of 0.26 (sd: 0.13) and nearly 100% call rate (sd: 0.05). On a chromosomal level, GBS-set missingness ranged from 9% to 15%, with a notable exception of the X-chromosome displaying over 30% missingness (Figure S5). Sample-wise genotype concordance between GBS and WGS data ranged from 82.6% to 97.5% (mean: 93.3%). A mere 1.3% of GBS-called homozygous variants were identified as heterozygous in the WGS dataset, and only 0.2% of heterozygous GBS variants were classified as homozygous in the WGS dataset. In total, 2 598 (12.5%) GBS variants were exclusive to the GBS call set, while 23 964 (56.8%) WGS variants were absent from the GBS (Table S2) variants.

Evaluating GBS based variant data for its ability to recover the realized relatedness matrix derived from >10 million bovine SNPs in the full genome data showed a convergence of both. With approximately 1 000 variants the matrices approach equivalence, as indicated by the eigenvalue distance dropping from >1 to approximately 0.15 (Figure 3). After this point, the GBS genotype-based matrices exhibited a slower convergence compared to the WGS-based counterpart. Results suggested that about 5 000 GBS markers equate to 2 000 WGS-derived SNP markers, fulfilling genomic selection needs.

**Figure 3.**
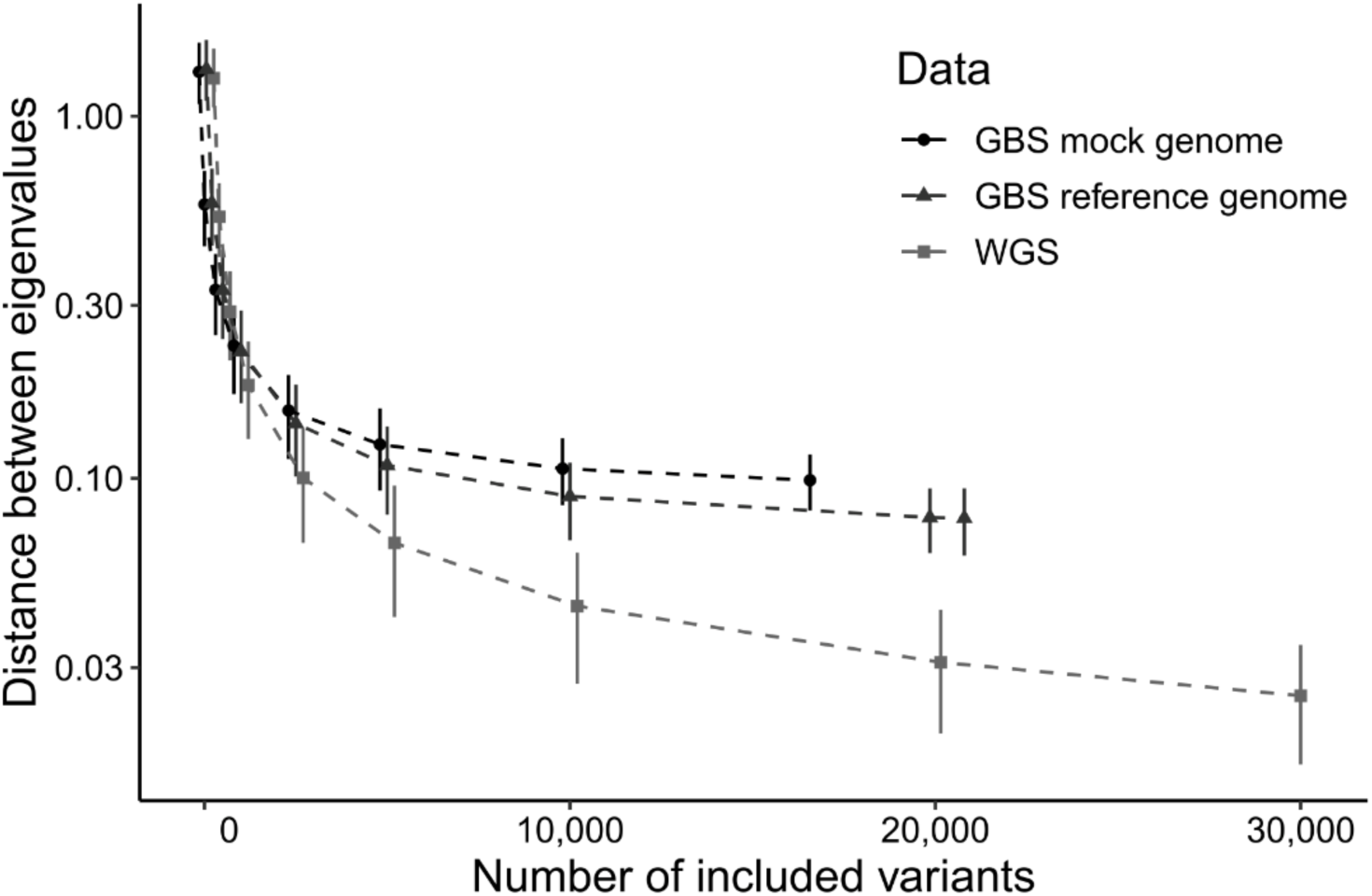
Evolution of eigenvalue distances as a function of the number of utilized DNA variants. The plot compares the distance between GRM matrix based on all whole genome sequence (WGS) derived variants and smaller variant subsamples from mock/reference GBS or WGS data. The plot displays the mean and 90% confidence intervals, generated from 1 000 bootstrapped resampling. Variant counts range from 50 to 30 000, encompassing the full GBS sample sets. The Y-axis is log-transformed to enhance visibility of differences.

### Proof of concept using non-model European whitefish species as an example

The European whitefish mock reference created by strategy 3, following the optimized mock creation parameters, was comprised of 159 403 clusters, spanning around 26 million bp, and suggested an average 4x – 8x fold read coverage. While shallow sequenced samples exhibited low coverage (4x), most samples demonstrated acceptable coverage (8x) against the created mock reference. Aligning the mock reference to the *Coregonus sp.* ‘balchen’ reference genome (AWG_v2) resulted in a coverage of 34 million bp due to multiple mapping, with alignment rates around 90% for quality-filtered PE reads against the mock reference and slightly higher (91%) against the AWG_v2 reference genome. Using an in-silico prediction for a 150-400 bp fragment size threshold led to 28 085 fragments and an approximate 80% alignment rate against this reference. Employing the mock reference facilitated calling 18 678 GBS variants, with a stable missingness below 5-7% for samples with over 1 million reads. Similarly, the existing reference genome enabled calling 23 275 GBS variants with a comparable stable missingness.

Genomic relatedness estimates between parent and offspring in whitefish trios averaged 0.53 (ranging 0.47 - 0.57) with the AWG_v2 reference genome data, and 0.49 (0.43-0.54) with the mock reference data aligning with the expectations [28]. Respectively, genomic relatedness among the parental fish averaged 0.09 ranging from −0.05 to 0.53 or averaging 0.08 and ranging from −0.04 to 0.49. Unrelated fish exclusively formed mated pairs (all relatedness estimates <0.05), aligning with expectations. Rare non-Mendelian inheritance, consistent across families, occurred in 3.3% (333.2 GBS variants on average) of the loci variable within the trios using AWG_v2 reference genome data and 3.4% (263.8 GBS variants on average) with mock reference data. Repeated Mendelian errors shared among loci were slightly smaller in the reference genome data (14.0%, 202 variants) compared to the mock reference data (14.8%, 167 variants). Both data sets exhibited similar estimates with a maximum absolute relatedness difference of 0.045 and generally agreed with prior pedigree knowledge.

### Repeatability

The repeatability assessment in bovine encompassed three separate runs: two utilizing 250 ng DNA (Orig- and RepI-set) and one employing 500 ng DNA (RepII-set) as starting material. All three sets underwent the same wet lab and optimized bioinformatic protocol using the ARS-UCD1.2 reference genome. The initial pipeline optimization run for the Orig-set yielded 20 794 GBS variants while the RepI-set and the RepII-set produced 19 066 and 19 988 GBS variants, respectively. Analyzing variant locations revealed a high degree of shared loci, with the RepI-set displaying 16 559 (79.6%) shared variants, and the RepII-set exhibiting 17 459 (84.0%) shared variants. Remarkably, the two repeated runs shared 16 556 variants in common, resulting in a cumulative sharing of 15 246 (73.3%) variants across all three runs (Figure 4a).

**Figure 4:**
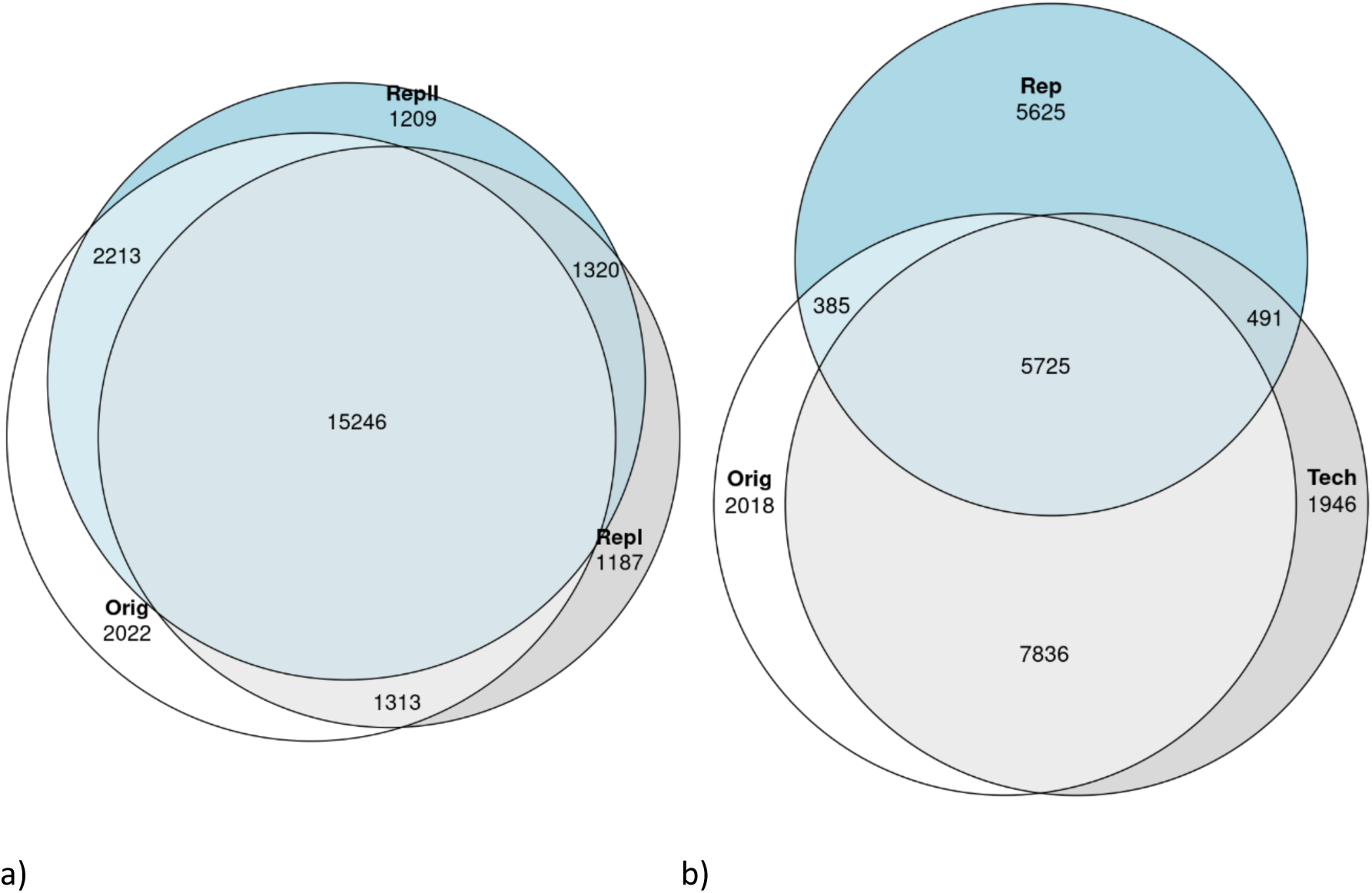
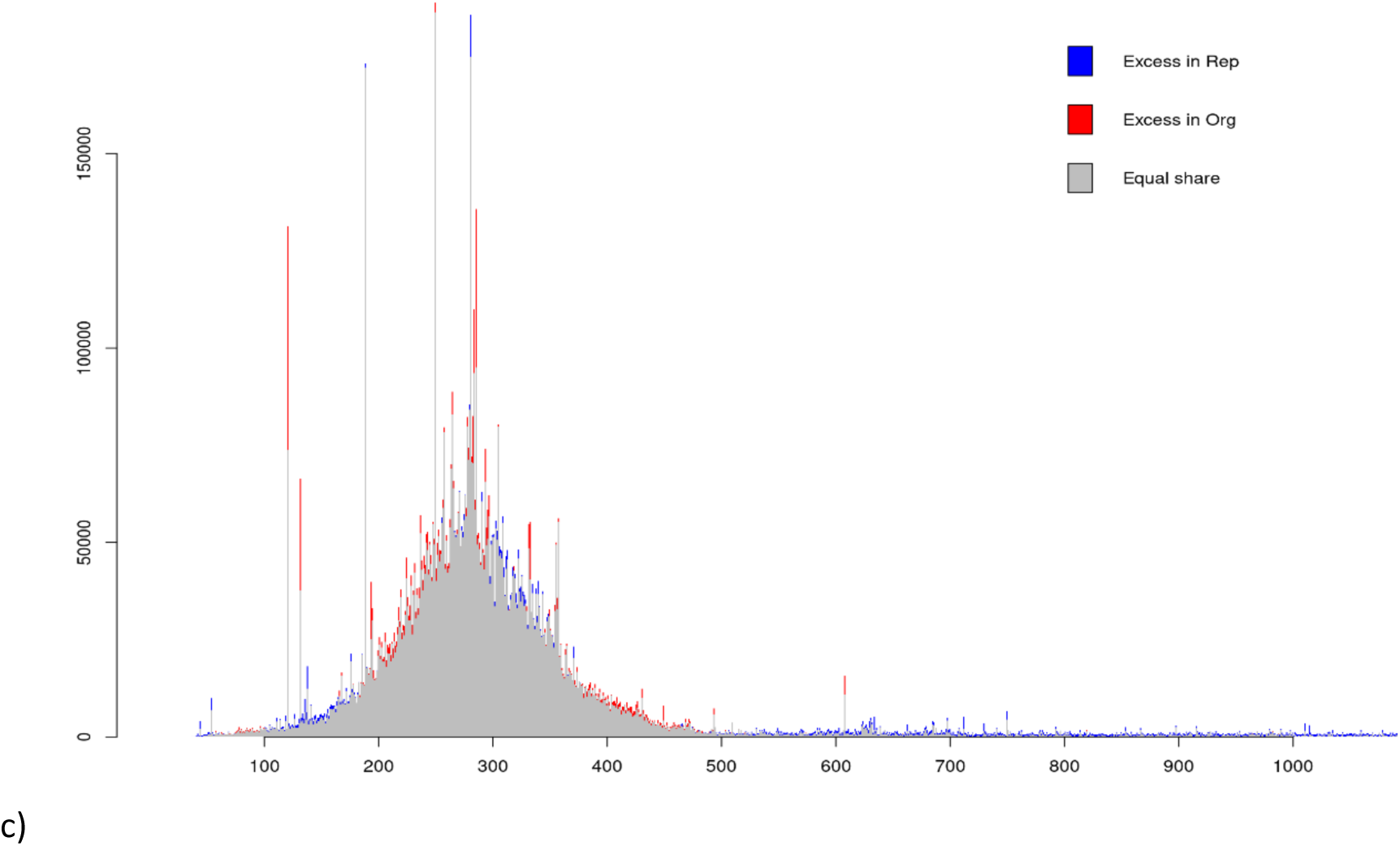
Repeatability intersection Venn diagram. Left side a) Cattle, right side b) Whitefish, c) read frequency distribution of the two Whitefish repeats.

Within the whitefish dataset, a repeatability analysis encompassing two distinct scenarios for a subset of 12 samples was performed. The first scenario involved technical replicates of identical libraries (Orig-set and Tech-set). In the second scenario, duplicate libraries were prepared from the same DNA samples (Rep-set). Dedicated pipeline runs for each set yielded 15 991 variants for the Orig-set, 16 025 variants for the Tech-set, and 12 253 variants for the Rep-set. Examination of intersecting variant locations highlighted a pronounced similarity between the Orig-set and Tech-set, sharing 13 561 (84.8%) loci. In contrast, the degree of sharing between the Orig-set and the Rep-set dropped to 6 110 (38.2%) and a similar value of 6 216 (38.8%) was observed for the Tech-set. Altogether, 5 725 variants were common to all three sets (Figure 4b). For the Orig-set as well as for the Rep-set the data aligned to the correct size selection range. However, the Rep-set had a slightly worse size range specificity but also less reads mapping to a few highly overrepresented sizes (Figure 4c).

Repeatability of individual variants at the whitefish sample level was also evaluated. For the 5 725 overall shared variants, 44.4% to 93.0% variants were equally called among repeated individuals. In pairwise comparisons, Orig-Tech samples shared 93.0% equally called variants, for the Orig-Rep comparison, however, on the average only 44.8% and in the Tech-Rep 44.4% of the variants were called equally. For the 15 246 shared variants across the three independently repeated cow GBS runs we obtained, however, for all three pairwise comparisons an average repeatability of over 90%.

Further, lift-over chains between the created mock references and the pre-existing reference genomes have been created to match variants called via the mock reference and those called by utilizing the pre-existing reference genome. For cattle, 16 571 variants were called using the mock reference. In total, 13 471 of these variants received successfully via lift-over a chromosomal location on the pre-existing reference genome. From these, 11 649 (>70%) intersected with the chromosomal location of variants called by utilizing the reference genome. In case of whitefish, from the 13 376 called variants via mock reference 10 693 could be lift-overed to the reference genome, with 6 481 (48.5%) variants having a chromosomal match with variants called based on the pre-existing reference genome.

## Discussion

We present here a GBS approach containing a refined ddRAD approach, where through the adaption of a published laboratory protocol [29] and the optimization and streamlining of the GBS sequencing data analysis steps utilizing the Snakemake workflow manager, we introduce a cost effective and robust genotyping procedure. RAD-Seq, since its inception by [17], has rapidly gained standing across diverse genetic research domains, spanning for example genetic map creation [14,30], mapping of production traits [31–33], population dynamics [34], and generating SNP resources for SNP array development [31,35]. Particularly, GBS stands out as a valuable tool for generating markers in non-model species with limited genome information. Our work extends the prior experimental demonstration of the ddRAD GBS method to facilitate genomic selection and breeding planning, especially for less studied farmed species. We successfully applied the developed protocol in non-model species (European whitefish), demonstrating its versatility and effectiveness, albeit revealing some remaining challenges.

The prevailing trend strongly favors incorporating bioinformatic workflow engines for robust pipeline implementations [36]. *Snakemake* [37], a widely adopted choice within the NGS field, was employed in our study to manage task dependencies, to reduce redundant computations upon pipeline re-execution, and to facilitate automated deployment, including integration with the *slurm* workload manager on our cluster. The native docker and singularity support enabled seamless utilization and versioning of necessary software tools. With a single command, the pipeline execution is initiated, channeling outputs into a well-organized main folder with structured subfolders housing the resultant analyses. This comprehensive strategy ensures full reproducibility and user-friendliness, accommodating those with limited programming skills, as all essential configurations are consolidated within a central configuration file. We chose *GBS-SNP-CROP* [38] as base solution as it utilizes the generated sequencing data in a straightforward way producing a large number of reliable variant genotypes [38]. We wrapped the well-established *GBS-SNP-CROP* pipeline into a *Snakemake* workflow and extended it with various steps to create an automatically generated report that allows the user to evaluate the GBS run and to trace possible problems with it.

### Data generation

We utilized the modified ddRAD method [29] for sequence data generation. By avoiding costly barcoded adapters and instead ligating digested fragments to non-barcoded adapters and utilizing standard Illumina dual-indexed barcodes for PCR enrichment and sample multiplexing, we reduced the library preparation costs to <9€/sample. While the laboratory workflow involves multiple steps that lack convenient commercial kits, optimization efforts streamlined the process. Hands-on-time was halved to 10 hr for 96 samples and 30 hr for 384 samples by normalizing DNA concentrations using Myra liquid handling system (Bio Molecular Systems, Australia), incorporating SPRIselect beads for size enrichment allowing to omit one of the two time consuming concentration measurements with Qubit. The utilization of BluePippin (Sage Science, USA) and other possible automations may further solidify routines and improve quality and time- and cost-efficiency.

By generating shorter 2×75 bp PE sequencing reads on the NextSeq550 we reduced sequencing cost to 10-14€/sample, with a yield of 1 million reads per sample. Utilizing shorter reads is advantageous over longer reads, as the aim is to use unlinked variants and to avoid the complications caused by closely linked markers in relatedness estimation [39]. Decreasing read length in favor of increasing the read depth helps in avoiding too low read depth, which may lead to under-calling the heterozygotes and incorrect assignment of them as a homozygotes [40]. Our results suggest that a sequencing depth exceeding one million reads per sample leads to a stable variant calling with minimal variant missingness in assessed species. However, the required sequencing depth highly depends on the number of targeted fragments, which is a balance between DNA quality, used enzymes, used fragment size range and the genome size of the investigated species and even the chosen sequencing technology. Moreover, the number of recovered variable sites depends on the genome variability. As a result, preliminary evaluation with a limited subset of samples is recommended to establish the balance between the targeted fragments and the minimum coverage threshold.

In European whitefish, around 40% of GBS variants were scored repeatedly across two fully independent analyses, aligning with earlier observations [29]. Conversely, in the bovine analysis, the first two repeats shared over 80% of the called variants, and all three repeats shared still approximately 75% of variants despite purposefully varying DNA amount. This indicated on the one hand a high level of repeatability achievable in certain species, and on the other hand, a remaining challenge in repeatability in other species. Here, e.g., a duplication [24] in the genome could cause read alignment issues that cannot be circumvented, and which could possibly cause differences in variant calling. In that case, filtering out paralogs as suggested by [30] could be a promising approach to follow.

General stochastic variability inherent in wet lab methods, encompassing fluctuations in PCR, library generation, and fragment size selection, plays a role in the repeatability [41]. These aspects may further interact with the applied bioinformatic methodologies. For example, DNA fragments carrying the reference allele are more likely to be successfully mapped or receive higher quality scores [42]. The repeatability is also influenced by the filtering steps during the variant calling phase, when various filters (MAF, minimum/maximum coverage as well as minimum call rate) are applied, as we confirmed comparing the pipeline reports for filtered and unfiltered variants (result not shown). Further, multi-mapping of reads might lead to unpredictable consequences. Notably even for European whitefish, repeated GBS variant scoring between technical replicates was frequent (85%), underscoring the potential enhancement of repeatability through simultaneous library preparation for all analyzed individuals, although the results suggested the non-repeating variants might partially represent repetitive genome segments. In cattle, where genomic selection relies on relatedness across generations, repeatability across fully independent analyses is of significance. Contrastingly, aquaculture-based genomic selection involves comparing reference populations and selection candidates within a generation [43], diminishing the need for repeatability across generations.

Additionally, relatedness estimation remains reasonably robust against missing data and genotyping errors when the variant count is substantial [22].

The GBS approach was tailored here for genomic selection utilizing a genomic relationship matrix, with the optimal informative GBS variant number falling between 1 000 and 10 000 [15] with a minimum of 1 000 – 2 000 SNPs generally suggested [15]. An in-silico comparison underscored the substantial influence of enzyme pair selection on reducing assessed genome complexity. However, even the enzyme pair with the lowest projected fragment count (EcoRI;SphI) was anticipated to yield ample variants. The difficulties of predicting fragment sequencing coverage are well-known and unassessed fragments are to be expected [41,44]. Accordingly, our final GBS variant numbers in cattle and whitefish (20k and 16k) reduced from their projections (36k and 21k forecasted). Unassessed fragments could arise from multiple factors, including genomic structural variations between references and samples, variation at restriction cut sites [45], and repeated regions, biased nucleotide content, and sequence length variation [41]. A sufficient variant number margin is preferrable, as breeders running a genomic selection program might prefer excluding low MAF variants increasing the variance of diagonal GRM elements [46] or variants with suspiciously high observed heterozygosity (>50%, [47]). Notwithstanding the challenges, the simple projections demonstrated to be sufficient for estimating variant number magnitudes for the ddRAD GBS method.

### Mock genome and pre-existing reference genome

For cattle a high-quality reference genome exists, while in our case representativeness of the European whitefish reference genome was uncertain. Utilizing a mock genome is essential when a reference genome is absent or incomplete for the target species [46,48]. Further, the spread between alignment rates for the existing reference genome and the created mock reference can serve as a metric for the evaluation of the representativeness of the reference genome for the data at hand. Acting as a stand-in scaffold or reference, the mock genome is essential for variant calling and the subsequent analyses by providing a foundation for aligning and mapping the sequencing reads as well as localizing the called variants. An effective strategy for determining cluster numbers include using either a small representative sample group or a single exemplary sample. The latter approach, however, may introduce biases from unique features of that single sample [46]. Constructing a mock genome from a broader sample range, although suggested [46], results in an inflated reference. Depending on the total number of samples and based on our observations, opting for a moderate collection of 3-5 samples minimize specific biases and avoids excessive inflation. The recommendation of Sabadin et al. [46], however, seems to be more relevant for heterogeneous sample sets, as they are common e.g., in plant breeding. In these cases, the introduced final mock correction step is expected to curb excessive cluster inflation. The refined final mock provides more stable results and is generally preferrable.

While a mock genome reference might be necessary, it is not curated against computational artifacts related to sequencing errors [49], sequencing or base composition bias [50–52], or repetitive regions [49] which can constitute 10-60% of the genome [53,54]. The suggested mock construction parameters are a good starting point for most animal species, but correctly separating duplicated genome regions while simultaneously collapsing and merging haplotypic differences into a haploid sequence is a challenge to all assemblers [55]. Here, we recommend several iterations of the pipeline with different settings especially for the identity criterion for merging clusters for each new GBS data generation case. The identity criterion can be increased until the alignment rate begins to decrease significantly while maintaining or increasing per-site coverage. Other parameters fine-tune the pipeline mainly by removing noise from the input data and have smaller impact. Given the influence of data and parameters on the created mock reference, archiving and sharing the reference facilitates later comparability and repeatability. Further, many pipeline parameters that had little impact in the present comparison, could get more influential for problematic data and as such could rescue still semi-optimal sequencing runs.

Using a subspecies-specific reference for cattle and a species group-specific reference for whitefish led to a 25% GBS variant increase over mock genomes, as expected when closely related reference genomes are available [47,56,57]. This underlines the advantage of employing reference genomes whenever feasible. While the surplus of variants might raise concerns about the genotype call quality, evaluating genotyping via Mendelian inheritance [58] contradicted this notion, showing stable and comparable inheritance error rates to reported NGS-generated SNP data [57–59]. Comparing GRMs between GBS and WGS sequencing favored the reference genome based GBS analysis, which approximated the WGS GRM matrix more closely. Despite the common concern of low MAF in GBS data [46], our comparison had lower MAF in the reference WGS data than in the GBS datasets. While the WGS data offers comprehensive insights, reference genomes are not flawless, for example, excluding variants on genome regions specific to individuals or populations [60,61] which may explain the minor difference between the two GBS GRM matrices. In general, using a very closely related reference genome increases the mapping and genotyping accuracy [56,62]. Therefore, it is recommended to execute both mock and possibly pre-existing reference genome paths of the pipeline and then compare the outcomes. Current observations suggest a reference genome is advantageous and should be used when available, though it is not an absolute requirement. Using a pre-existing reference genome offers a high quality assembly and consistency and possibly annotated genomic context for interpretation [63]. Further, the use of a reference genome facilitates evaluating the representativeness of the data and allows linkage-based analyses.

Variant calling using different mock genomes or a pre-existing reference genome might include different variants [38], but the approaches gave currently very similar relatedness estimates. This aligns with previous studies suggesting that while extensive repeatability of GBS genotype data can be challenging biological inferences based on these data sets are more robust [20,64,65]. When genomic selection analyses are based on relatedness, fixing the reference genome is not the only option for merging data sets, since it is possible to combine partially overlapping relatedness matrices [66]. However, this necessitates having representative population samples with reference individuals of varying relatedness for both having reliable estimates within each round and for enabling merging of the matrices. Comparability issues might occur even when basing analysis on reference genomes, which develop over time [67].

## Conclusions

The relatedness estimates based on the developed ddRAD GBS protocol aligns with independent relatedness estimates in both cattle and European whitefish samples, showcasing its versatility and extending the performance demonstration beyond GBS-SNP-CROPs original aim of identifying biological replicates. Our results conclude that while a pre-existing reference genome enhances variant calling quality and quantity, its absence does not impede the GBS-based genomic evaluation or selection. The applicability of the presented approach for genomic evaluation has been demonstrated for European whitefish [68], despite its challenging genomic structure. Further optimization, including fragment size window refinements and incorporation of methylation-sensitive restriction enzymes [69] could bring even greater efficiency and accuracy. The robust and user-friendly bioinformatic pipeline with an implementation of best practice approaches and wet-lab workflow achieves our broader goal of democratizing genotyping methods for researchers with varying levels of bioinformatics expertise and across a wide range of species and especially in less-studied production species. Experimenting with individual tuning parameters for the data at hand remains, however, indispensable and normally several pipeline runs are required until satisfying results are obtained. Furthermore, adjusting the filtering thresholds of called variants according to the analysis scope is still a required step, though default values should work well in many situations.

## Methods

### Samples

Altogether 12 Nordic Red dairy cows from the Luke research barn were selected for GBS and WGS sequencing, pipeline optimization and benchmarking. For each cow sample three repeated GBS libraries and one WGS library were created, starting from the same extracted DNA so that in total 36 GBS libraries and 12 WGS of cow samples were sequenced (Figure S1).

In addition, 42 European whitefish were used for pipeline validation and repeatability testing. Fish samples consisted of 27 randomly picked, unrelated individuals and 5 families of trios (parents and one offspring). From the set of random individuals, 12 whitefish were sequenced three times, twice with technical replicates of the same library and once with an entirely new library, that was started from the DNA. The European whitefish originate from the national breeding program maintained by Luke at the inland, freshwater fish farm located in Enonkoski [25,26]. The broodstock was established in 1998 from an anadromous wild strain of the river Kokemäki. Currently, the breeding program is based on traditional sire-dam-offspring pedigree, maintained by the use of family tanks during the early phase of growth [25,26], but the development of SNPs will enable to implement also genomic selection.

Cow DNA was extracted from blood (ethical permission ESAVI/16348/2019) while fin tissue preserved in 100% ethanol was used for DNA extraction from fish. DNA was extracted using DNeasy Blood & Tissue Kit (Qiagen, Germany) following manufacturer’s protocol.

### Enzyme selection in silico

Restriction enzyme pairs for genome reduction were selected i) to generate a number of fragments providing above 5 000 GBS variants and ii) to leave a suitable overhang for library preparation. Assuming the proportion of variable sites of approximately 0.005 [24] and aiming for Paired-End (PE) sequencing with a total of 150 (2×75 bp) sequence read length per fragment, the number of variable sites was expected to be 0.75 times the fragment number. That suggested inclusion of at least 10 000 fragments, if all variable sites pass all quality ascertainment steps. The considered restriction enzyme pairs were EcoRI with MspI, SphI, MseI and NlaIII, or SphI with MluCI. These enzymes were previously used successfully for GBS in other species [21,70,71]. For a wider applicability, six reference genomes were included for the restriction enzyme evaluation: *Bos taurus* (ARS-UCD1.2), *Coregonus supersum ‘balchen’* (AWG_v2), *Gallus gallus* (GRCg6a), *Hermetia illucens* (iHerIll2.2), *Oncorhynchus mykiss* (Omyk_1.0), and *Salmo salar* (ICSASG_v2). DdRAD library construction was simulated using SimRAD version 0.96 [72], but the functions were adjusted to use the full cut site. Digestion was simulated by using both the full reference genome contigs as well as reduced genomes of 10 random 10% genome subsamples. The full genome based (*Bos taurus* and *Coregonus supersum*) predicted fragments for the chosen EcoRI;SphI enzyme pair were used for quality evaluation of the GBS analysis. The obtained sequence data was used to estimate the effective size window and as consequence the size selection window was set to 150 - 400, for consistency. The effective size window thresholds were roughly estimated as values, where the slope of the density curves of the aligned fragments turned to +1 (lower size threshold) and −1 (upper size threshold).

### ddRAD library preparation

The workflow (Figure S6) for the ddRAD library preparation was adapted from [29]. In detail, 250 or 500 ng of DNA was double-digested with two restriction enzymes EcoRI-HF (G^AATTC) and SphI-HF (GCATG^C) (New England Biolab, USA). The restriction reaction was performed in a volume of 20 µL, containing 17 µL of DNA (250 ng/500 ng in total), 0.25 µL of EcoRI-HF (5 units), 0.25 µL of SphI-HF (5 units), 2 µL of cut-smart buffer (10x) and 0.5 µL of molecular grade water at 37°C for 2h, following heat-inactivation for 15 min at 65°C. Two non-barcoded restriction site specific adapters (Table S3) were ligated by adding 1 µL of each adapter (adapter P1 EcoRI: 1 µM, adapter P2, SphI: 10 µM) to the restriction mixture, 0.5 µL of T4 ligase (200 units) and 1.5 µL of ligation buffer (New England Biolab, USA). Ligation was performed at 16°C for 14h, following heat-inactivation at 65°C for 15 min. DNA-fragments were selected between 200 bp and 700 bp by using SPRIselect magnetic beads (Beckman Coulter, USA) with a left-right ratio of 1x-0.56x. In details, the volume of each sample was adjusted with molecular grade water to 50 µL and then 28 µL of SPRIselect beads were added to achieve a 0.56x ratio for the selection of fragments shorter than 700 bp following selection of fragments longer than 200 bp by adding 22 µL of SPRIselect beads to achieve a ratio of 1x. The size selected DNA was resuspended in 25 µL of molecular grade water. Samples were barcoded by adding Illumina Nextera v2 (Illumina, San Diego, CA, USA) combinatorial dual-indexed barcodes (i7 and i5). For each individual sample a PCR-mix containing 6 µL of 5x Phusion HF buffer, 0.4 µL dNTP (10 mM), 0.2 µL of Phusion HF DNA polymerase (0.4 units) (ThermoFisher scientific, USA), 1.5 µL of i5 barcode primer, 1.5 µL of i7 barcode primer, 5 µL of sample and 15.4 µL of molecular grade water was prepared, two PCR reactions per sample were performed. The cycling conditions were as follows: initial denaturation at 98°C for 30 sec, followed by 18 cycles of 10 sec at 98°C, 20 sec at 61°C, 15 sec at 72°C and a final elongation step at 72°C for 10 min. The two PCR reactions per sample were pooled, the volume was adjusted to 50 µL, and small fragment removal was carried out with 40 µL (0.8x) SPRIselect beads. The size selected PCR products were resuspended in 25 µL molecular grade water and quantified using Qubit Flex with 1x dsDNA HS assay (ThermoFisher scientific, USA). Only products with a significantly higher amount than the No Template Control (NTC) were used for sequencing (>3 ng/µL).

### Sequencing

Single ddRAD libraries were pooled in equimolar amounts. The pool was size selected with SPRIselect beads to the length between 300 and 700 bp (ratio 0.75-0.56x), corresponding to the combined length of 150-550 bp restriction insert and 147 bp adapter. The quality and size of the pooled sequencing library was evaluated on the TapeStation 4150 (Agilent, USA) using the DNA HS1000 assay.

Quantification of the library was done using Qubit 4 (1x dsDNA HS assay) (ThermoFisher scientific, USA). Following the guidelines from the NextSeq System denature and dilute libraries guide (Document # 15048776 v09, December 2018 (Illumina, San Diego, CA, USA)), the library was diluted for sequencing to a final concentration of 1.4 pM, containing 10% PhiX control, to increase complexity at the start of the sequencing. The PE sequencing (2×75 bp) was performed on the NextSeq 550 (Illumina, San Diego, CA, USA) using medium output flow cell.

The WGS of cow samples was performed at the Finnish Functional Genomics Centre (Turku, Finland) using TruSeq® DNA PCR-Free Library kit (Illumina, San Diego, CA, USA) and PE sequencing (2×150 bp) on an Illumina NovaSeq 6000 (Illumina, San Diego, CA, USA) platform.

### Mock-reference genome

Analyzing GBS data without a preexisting reference genome necessitates in creating a technical (mock) reference. For this, various sample selection methods were considered: choosing the sample with the highest read count (mock-strategy 1), a sample with an average read count (mock-strategy 2), a random subset of three samples (mock-strategy 3), or all samples (mock-strategy 4).

As the first step, the raw PE sequences were checked for overlap that might happen in case of short inserts. Overlapping reads were merged into single-end (SE) reads using *PEAR* [73], with two tuning parameters being optimized here: the *p* option (values between 0.001 and 0.1) for a statistical test to determine read-pair merging, and the *pl* option (values 30 to 70) for defining the minimum accepted total length of the merged construct. These parameters determined when read pairs were merged and whether the construct’s length met the criteria for inclusion. PE reads that could not be merged, were then stitched together with a sequence of 20 N bases as standard for the pipeline. Stitching of reads was controlled by the parameter *rl*, and reads were stitched, if the length of read1 was larger than (rl - 19) and length of read2 was larger than (rl - 5), otherwise reads were not used for the mock generation. The resulting SE reads were utilized to construct the de-novo mock reference genome using *vsearch* [74]. In the de-novo building phase, two *vsearch* options were fine-tuned: the *id* option (values between 0.8 and 0.99), defining the minimum pairwise identity for merging two clusters, and the *min* option (values between 80 and 160), setting the minimum cluster length for inclusion in the mock reference. The in-silico simulated protocol as described in “Enzyme selection in silico” was used to evaluate the mock reference constructs.

Following the de-novo mock reference creation, an additional refinement step was applied, where clusters with low coverage were removed from the mock reference. Tuning parameters were *totalReadCoverage* and *minSampleCoverage*. The first parameter defines the minimum number of reads that need to be aligned across all samples on a cluster to keep it in the mock reference. The second parameter defines the minimum number of samples that need to have at least a single read aligned to a cluster so that this cluster remains in the mock. For the tuning of the *totalReadCoverage* we tested 6, 12, 24, 60 and 120 as values and for *minSampleCoverage* reads from 2 (10%), 4 (25%), 6 (50%), 8 (75%), 10 (90%), 12 (100%) of the total number of samples in the study.

### Variant calling

The GBS variant calling was done using *Snakebite-GBS* [75], which is a *Snakemake* pipeline extension that is based on the existing *GBS-SNP-CROP* [38] pipeline and that is part of the Snakebite framework *Snakepit* [76]. First, the quality-trimmed reads were aligned with *BWA-mem* [77] against the mock and/or preexisting reference genome(s). Then, *samtools mpileup* [78] was used for variant calling and various filters were applied to obtain the final variant set. The underlying GBS-SNP-CROP pipeline allows for eight different filters: (1) *mnHoDepth0* (value: 5), the minimum depth required for calling a homozygote when the alternative allele depth equals 0; (2) *mnHoDepth1* (value: 20) the minimum depth required for calling a homozygote when the alternative allele depth equals 1; (3) *mnHetDepth* (value: 3) the minimum depth required for each allele when calling a heterozygote; (4) *altStrength* (value: 0.8) the minimum proportion of non-primary allele reads that are the secondary allele; (5) *mnAlleleRatio* (value: 0.25) the minimum required ratio of the less frequent allele depth to the more frequent allele depth; (6) *mnCall* (value: 0.75) the minimum acceptable proportion of genotyped individuals to retain a variant; (7) *mnAvgDepth* (value: 3) the minimum average read depth of an acceptable variant; (8) *mxAvgDepth* (value: 200) the maximum average read depth of an acceptable variant.

The cattle WGS variant calling was performed following the *GATK4* best practices [79] implemented as *Snakemake* [37] workflow called *Snakebite-WGS* [80]. Implemented steps contain, among others, the GATK base recalibrator as well as a model to adjust the base quality scores and a base recalibration step. Variant calling is done via haplotype caller. The pipeline utilizes also BWA-mem to align the data but includes a refinement step using *Picard* before the *GATK4* software suite is used for the final variant calling with applied default filters.

### GBS quality evaluation

The generated cow GBS variant data was mapped against an in-silico digested ARS-UCD1.2 reference genome for evaluating the size selection performance. Following variant calling, sample-wise genotype concordance between GBS and WGS sequencing strategies was assessed using *Picard*.

The repeatability of the GBS runs was tested by intersecting the variant locations on the corresponding reference genomes. Here, *bcftools* [81] was used to intersect the three vcf-files and corresponding intersection numbers were calculated. Further, *samtools mpileup* was run for the GBS data aligned to the reference genome and for each sample contiguous areas, that had a minimum coverage of three reads, were identified and stored in bed-format. Individual sample-wise bed-files were then merged and only regions with read support from at least 10 samples were kept. This bed-file was then used to intersect the WGS-based vcf file using *bedtools* [82] and extract WGS variants only from the corresponding intersecting genome regions.

In cattle, the GBS variant based variability and relatedness were compared against resampled WGS variants with restricted variant numbers from 50 to 30 000 to compare how the variant number influenced the classical Genomic Relatedness Matrix (GRM) calculated using the R-package *BGData* [83]. The GRM based on the full WGS variant matrix was compared to smaller bootstrap samples of WGS and GBS data.

The lift-over between mock reference and pre-existing reference genome to compare variants from both methods based on their chromosomal was done by using the tool *transanno*. Here, first the mock reference was aligned against the reference genome and the resulting file in pairwise mapping format (paf) was then used in *transanno* to create the lift-over chain and eventually to perform the lift-over. Chromosomal locations between the lift-overed mock reference-based variants and their pre-existing reference genome based counterparts were then again matched via *bcftools isec*.

The GRM structure differences were quantified by measuring the variability in different directions using the distance between the eigenvalues of the matrices, calculated using the Frobenius matrix norm.

For whitefish data, relatedness in trios was assessed using the full whitefish data set to overcome bias in the small data set caused by few closely related individuals in the parental generation. In addition to the genomic relatedness, the genotype quality was assessed by evaluating non-Mendelian inheritance of the GBS variants in five families of trios, that included parents and an offspring.

## List of abbreviations

Bp: basepair
ddRAD: double-digest RAD-sequencing
GBS: Genotyping-by-sequencing
GRM: Genomic Relatedness Matrix
MAF: Minor Allele Frequencies
NGS: next generation sequencing
PE: Paired-end
RAD: Restriction-site associated DNA sequencing
SE: Single-end
SNP: Single Nucleotide Polymorphism
WGS: Whole-Genome-Sequencing

## Supplementary Information

Additional file 1.docx : Contains all referred Supplemental Figures S1-8 Additional file 2.xlsx: Contains all referred Supplemental Tables S1-3

## Declarations

### Ethics approval and consent to participate

The study was performed in accordance with Finnish animal welfare legislation and complied with the directive 2010/63/EU implemented in Finnish legislation in the Act on the Use of Animals for Experimental Purposes (62/2006). All experimental fish were anaesthetized with tricaine methanesulfonate before sampling to minimize suffering. Cattle: ethical permission ESAVI/16348/2019)

### Consent for publication

Not applicable.

### Availability of data and materials

The datasets generated and analyzed during the current study are available in the European Nucleotide Archive (ENA), accession number PRJEB66491.

## Author contributions

AK, IT, MT, TIT: conception of the study; IT, AK: funding acquisition; OB: laboratory work; DF, MT: data analysis and writing the manuscript; IT, TIT, OB, AK: manuscript revision. All authors approved the final manuscript.

## Acknowledgements

We thank Finnish Functional Genomics Centre, supported by University of Turku, Åbo Akademi University and Biocenter Finland, for whole genome sequencing services and CSC – IT Center for Science, Finland, for computational resources. Antti Nousiainen and Heikki Koskinen are acknowledged for providing whitefish samples.

