## Supplemental Figures for "Fine-Tuning GBS Data with Comparison of Reference and Mock Genome Approaches for Advancing Genomic Selection in Less Studied Farmed Species"

### Supplemental material

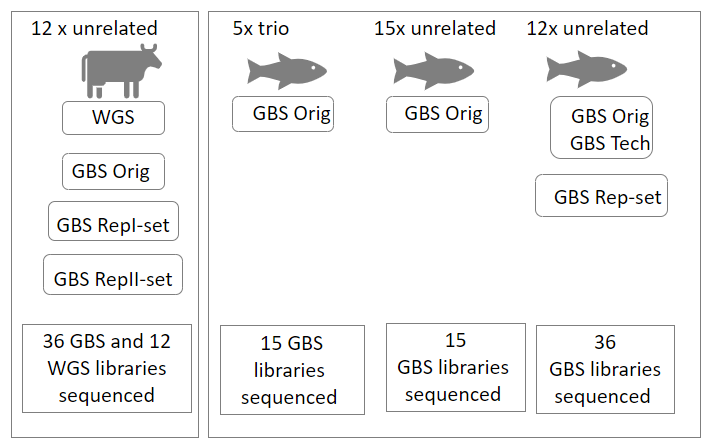

Figure S1. The schematic representation of the sequencing strategy. Altogether 12 cattle and 42 European whitefish individuals were used to create 36 cow and 66 whitefish GBS datasets. GBS Orig = The GBS dataset produced starting from DNA, GBS Rep-set = The GBS dataset produced from a new sequencing library, created from the same DNA extract as in GBS Orig dataset, GBS Tech = The GBS dataset produced by re-sequencing aliquots of the same sequencing library prepared for GBS Orig dataset

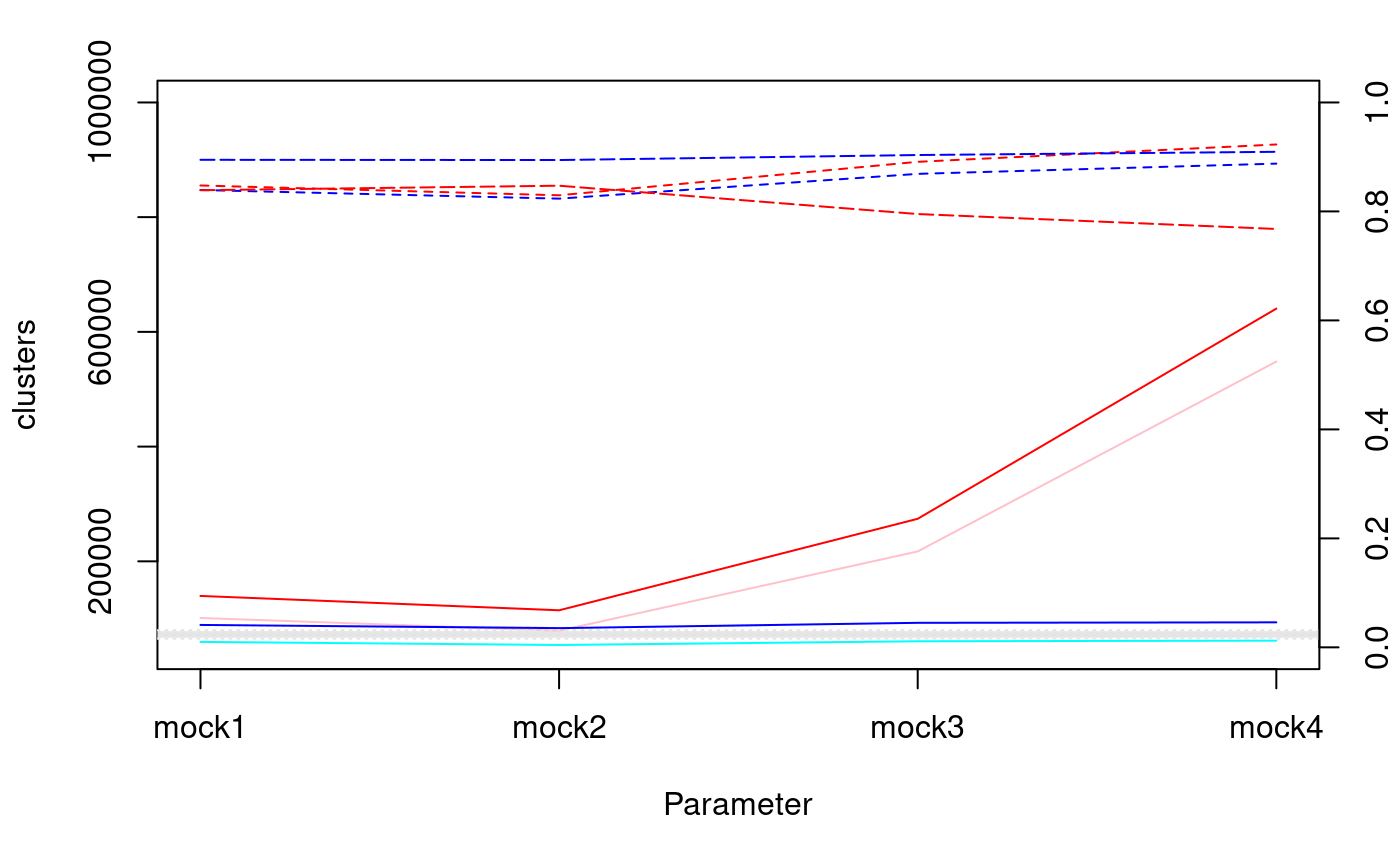

Figure S2: Quality assessment metrics for the alignment of different mock sample sampling strategies. The gray box indicates the aimed target, based in the in-silico prediction. Mock 1: sample with the highest read count; mock2: sample with an average read count; mock3: random subset of three samples; mock4: all samples.

Solid lines refer to left axis (clusters), dashed lines to right axis (ratios). Redish colours indicate the mock reference, blueish lines the refined mock reference. Intense colours (red, blue) indicate the number of aligned clusters from the mock reference to the reference genome, light colours (pink, cyan) the number of secondary alignments of that alignment.

Dashed lines indicate the average alignment rates across the samples against the mock (red) and refined mock (blue), long-dashed lines indicate the alignment rates of the mock reference (red) and refined-mock reference against the in-silico predicted reference genome.

The grey box indicates the predicted number of clusters after digestion.

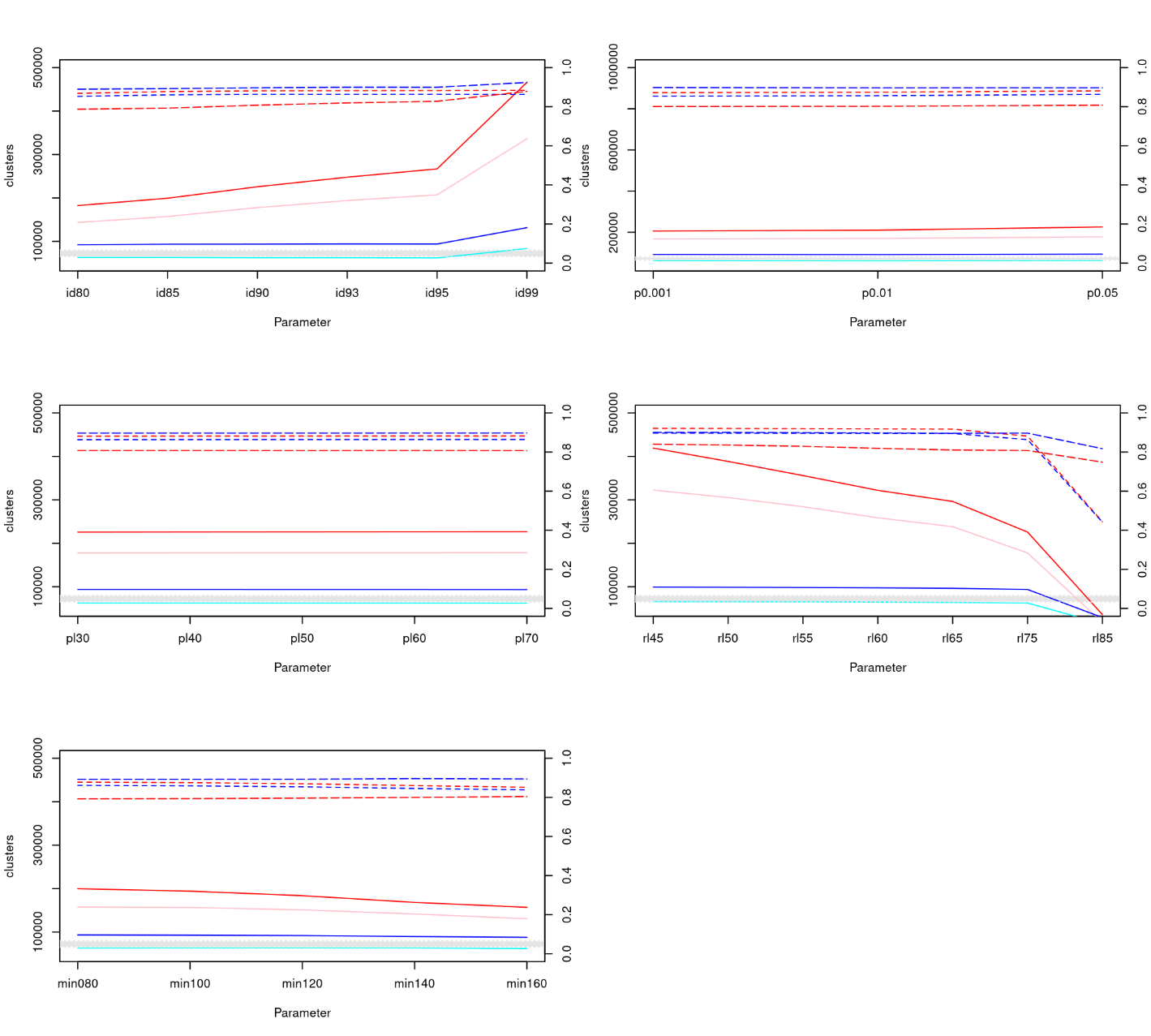

Figure S3: Impact of different parameter choices on various pipeline tuning parameters. Solid lines refer to left axis (clusters), dashed lines to right axis (ratios). Redish colours indicate the mock reference, blueish lines the refined mock reference. Intense colours (red, blue) indicate the number of aligned clusters from the mock reference to the reference genome, light colours (pink, cyan) the number of secondary alignments of that alignment.

Dashed lines indicate the average alignment rates across the samples against the mock (red) and refined mock (blue), long-dashed lines indicate the alignment rates of the mock reference (red) and refined-mock reference against the in-silico predicted reference genome.

The grey box indicates the predicted number of clusters after digestion.

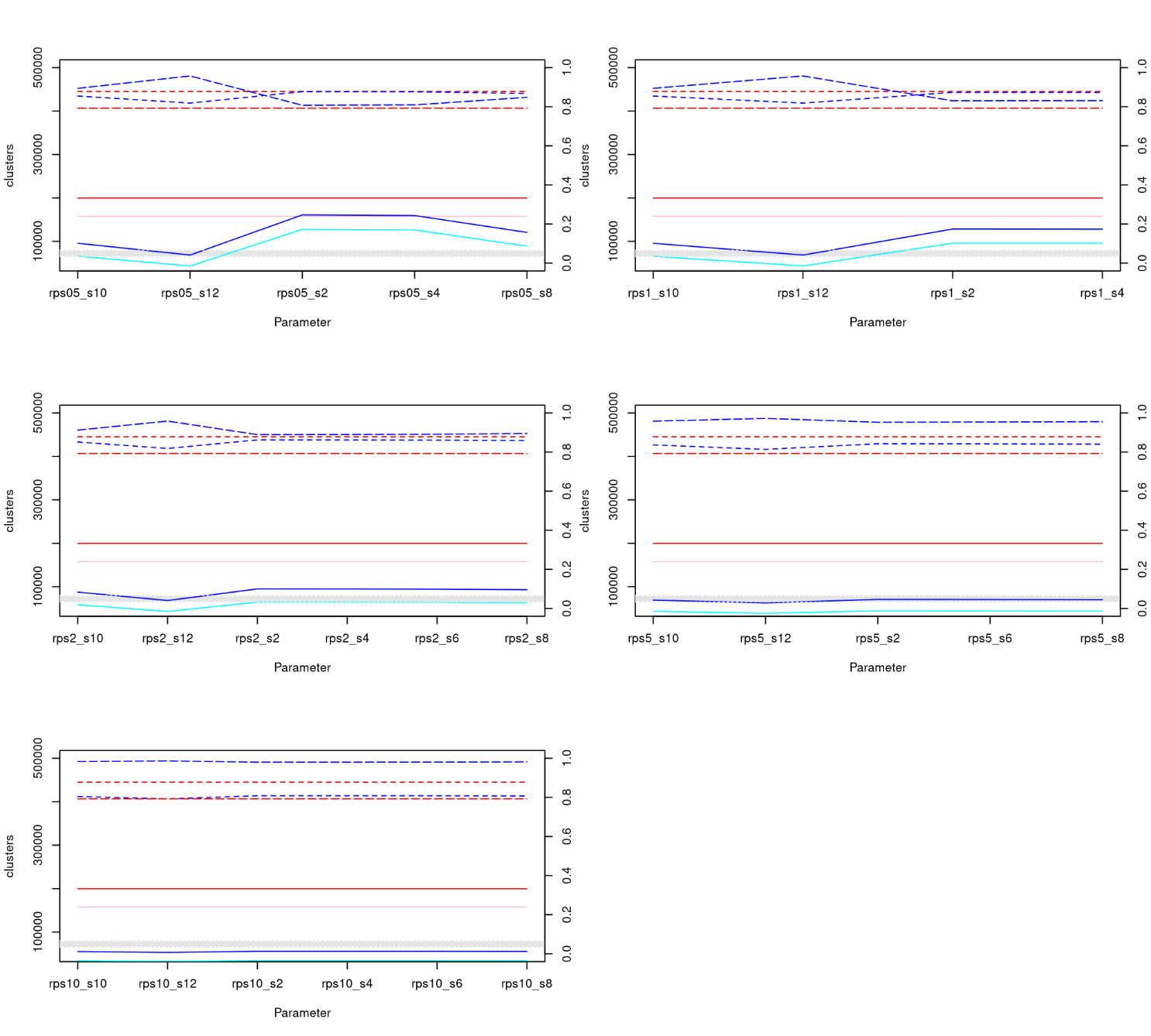

Figure S4: Impact of tuning parameters for the final mock. Tested values are here average number of reads per sample (rps) and number samples with reads on cluster (s)

Solid lines refer to left axis (clusters), dashed lines to right axis (ratios). Redish colours indicate the mock reference, blueish lines the refined mock reference. Intense colours (red, blue) indicate the number of aligned clusters from the mock reference to the reference genome, light colours (pink, cyan) the number of secondary alignments of that alignment.

Dashed lines indicate the average alignment rates across the samples against the mock (red) and refined mock (blue), long-dashed lines indicate the alignment rates of the mock reference (red) and refined-mock reference against the in-silico predicted reference genome.

The grey box indicates the predicted number of clusters after digestion.

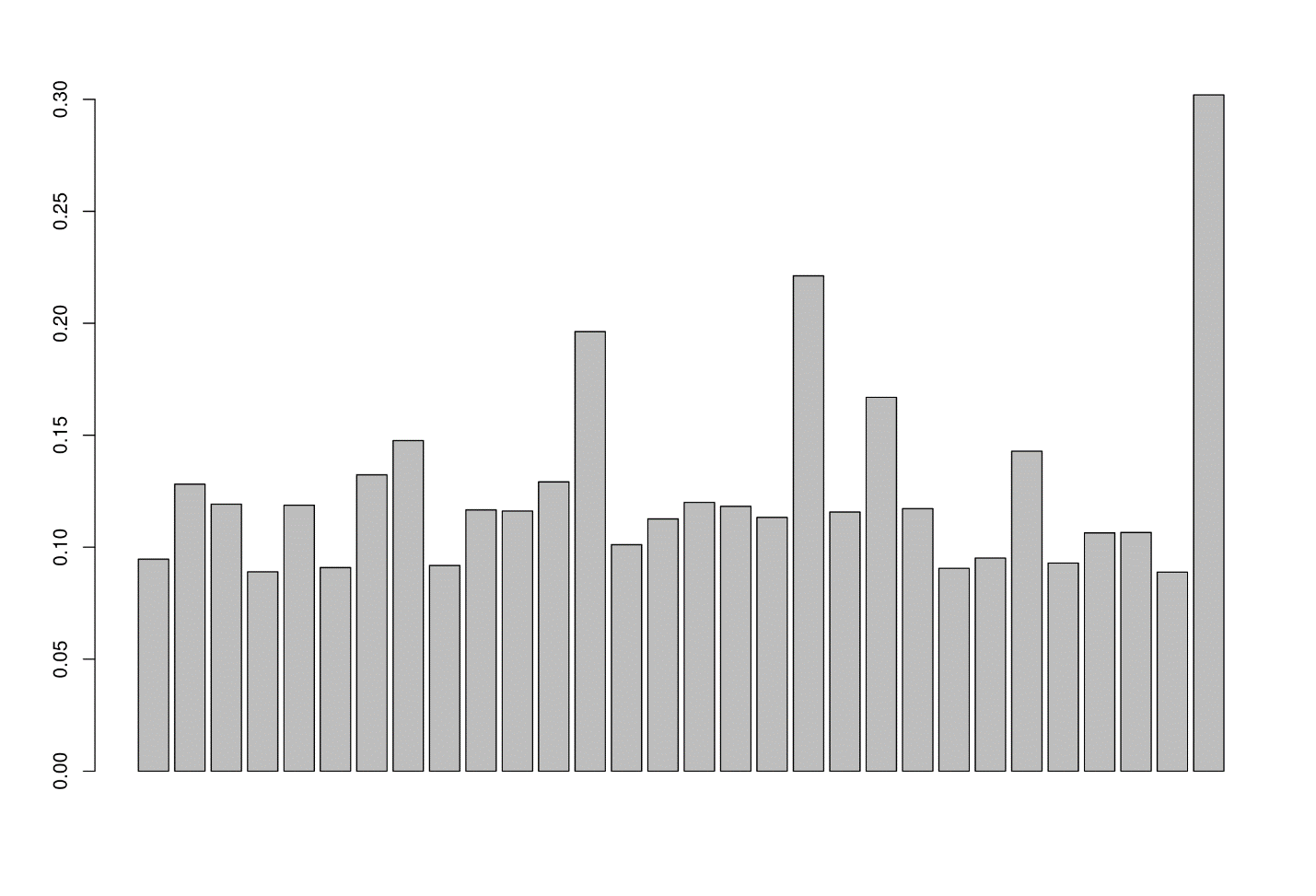

Figure S5. Ratio of variants that could not be recovered from the WGS data, split per chromosome 1-29+X.

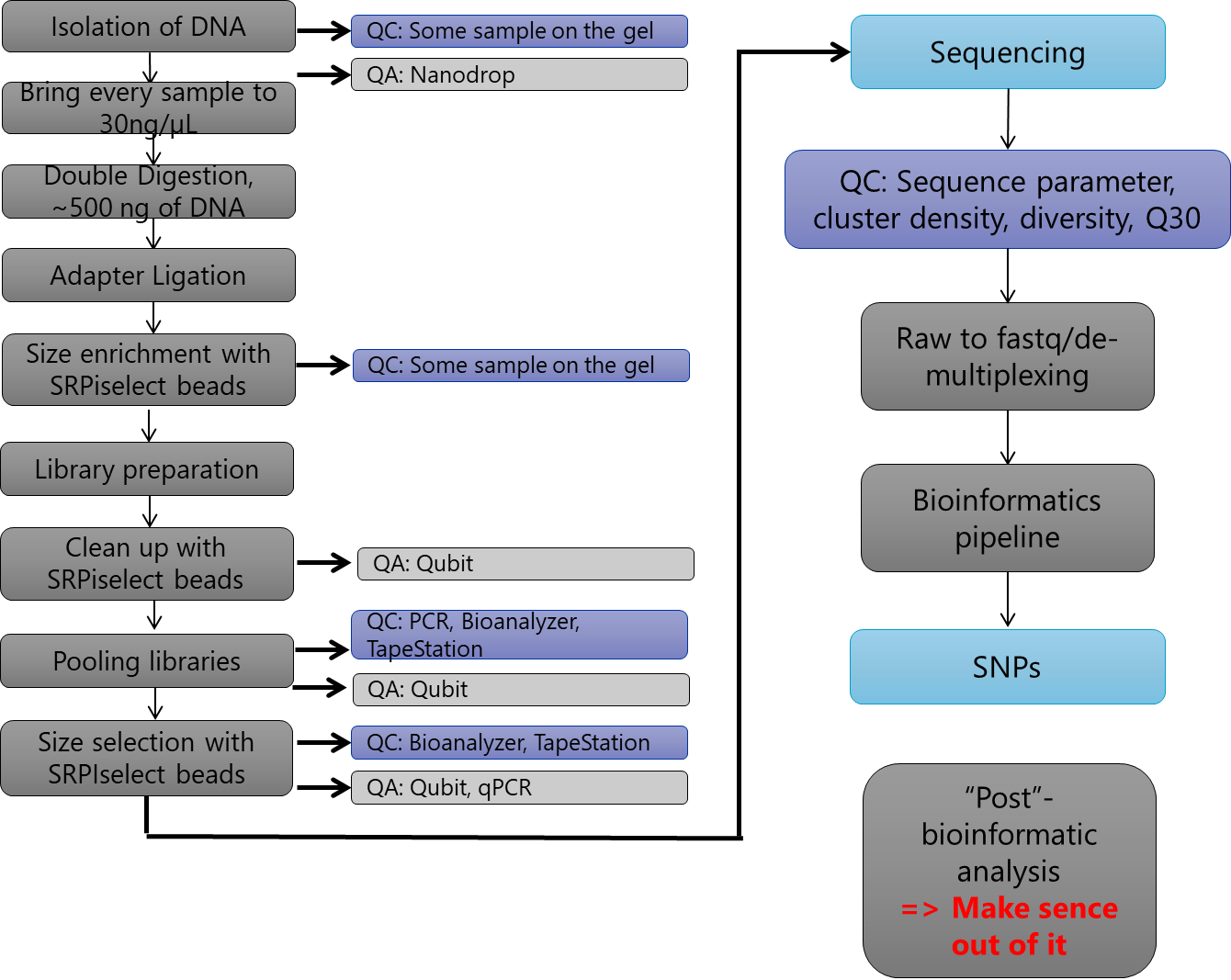

Figure S6: Workflow diagram of the wetlab pipeline

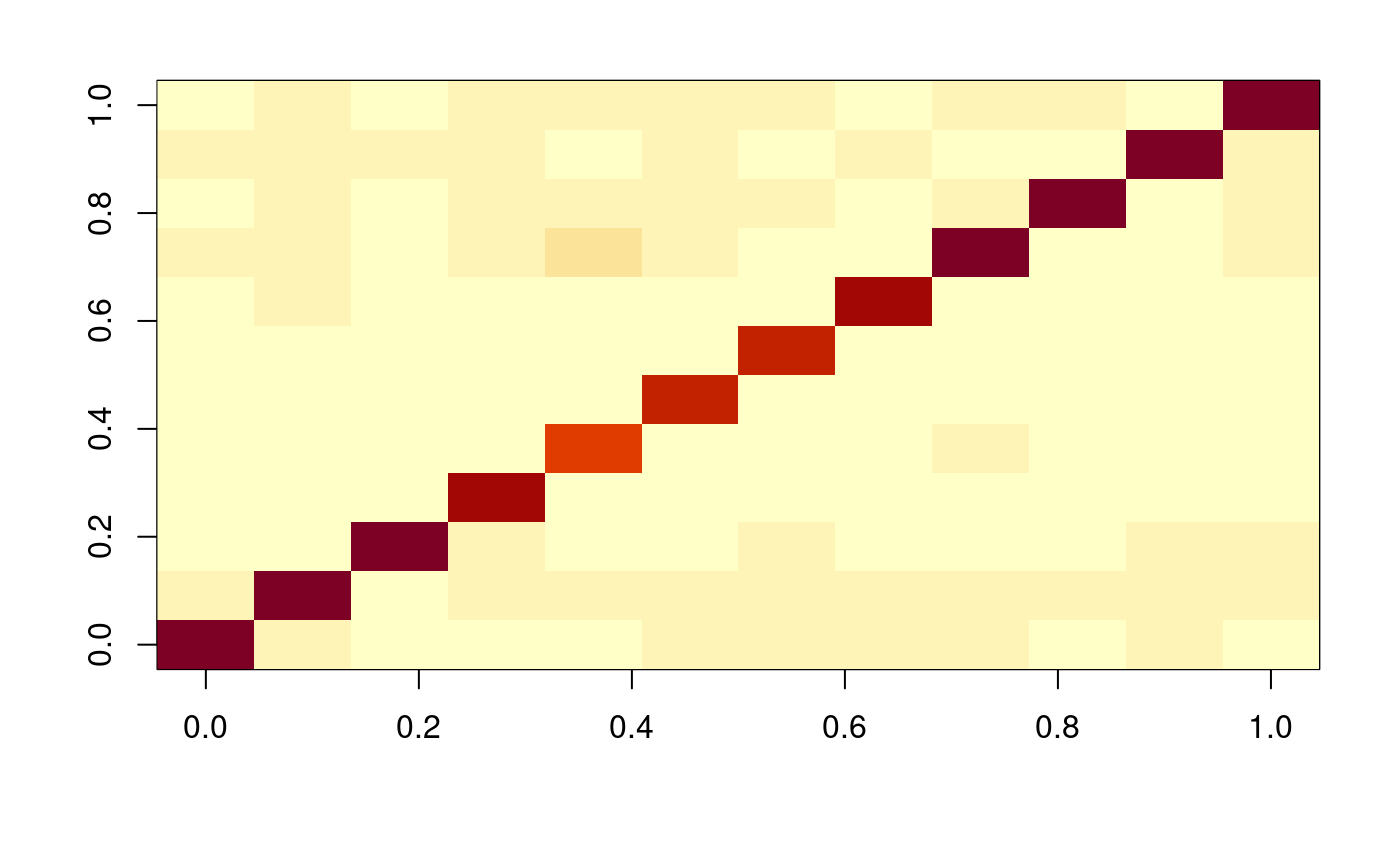
Figure S7: Heatmap of the pairwise genotype concordance between samples. It clearly shows that no sample mixup happened and that the two pipelines agree very well.
